# Combination antagonism of TNF superfamily signaling for T cell immunosuppression

**DOI:** 10.64898/2026.04.27.721101

**Authors:** Praveen Krishna Veerasubramanian, Wenlan Zang, Vijaya Amancha, Thomas A. Wynn, Jie Quan, Fridrik J. Karlsson

**Author notes:** Equal contributors (Jie Quan, Fridrik J. Karlsson). Corresponding author, Inflammation and Immunology Research Unit, Pfizer Research and Development, 1 Portland Street, Cambridge, MA 02139.

## Abstract

The tumor necrosis factor (TNF) and TNF receptor (TNFR) superfamilies comprise 47 proteins that regulate immune signaling and T cell costimulation. While TNF inhibitors are established therapies for immune-mediated inflammatory diseases (IMIDs), their efficacy is limited by primary non-response and loss of efficacy over time. Preclinical evidence suggests that TNF/TNFR members exhibit redundant and synergistic signaling, motivating combination targeting strategies. In this study, we have systematically evaluated TNF/TNFR combinations as potential immunotolerance targets using integrated computational and experimental approaches. We applied a gene prioritization framework incorporating transcriptomics, genetics, druggability, and pathway regulation data to derive disease association scores for the TNF/TNFR genes in rheumatoid arthritis and inflammatory bowel diseases. These scores, together with T cell expression profiling, were used to prioritize ten targets for combinatorial screening in mixed lymphocyte reactions using clinical-stage and preclinical pharmacological inhibitors. Four combinations of drugs inhibiting TNF+CD40L, TNF+OX40L, CD40L+OX40L, and CD40L+LTβ/LIGHT each in combination significantly reduced T cell production of IL-2 and IFN-γ. RNA-Seq analysis revealed that these combinations downregulated genes involved in T cell activation, proliferation, differentiation, and cytokine production that were upregulated during allogeneic responses. Notably, TNF+CD40L co-inhibition (Adalimumab+Dapirolizumab) produced the most robust suppression, uniquely downregulating 337 genes enriched for T cell activation pathways including NF-κB, ERK1/2, and cytokine production. These findings demonstrate that combinatorial TNF/TNFR targeting can potently suppress allogeneic T cell responses and support further preclinical evaluation as a tolerance-inducing therapeutic strategy for refractory IMIDs.

## Introduction

The tumor necrosis factor (TNF) superfamilies of ligands and receptors (TNFR) comprise 47 proteins that play essential roles in immune signaling and homeostasis. Among them, the eponymous TNF (previously TNF-α) is the most extensively studied member, known for its systemic inflammatory effects that characterize several immune-mediated inflammatory diseases (IMIDs). TNF inhibitory biologics (TNFi) have been a cornerstone of IMID treatment since their clinical introduction in the early 2000s. The broader TNF/TNFR families include key co-stimulatory molecules including OX40, 4-1BB, GITR, CD27, CD30, and HVEM, which fine-tune T cell responses. Dysregulation of these signaling pathways is implicated in multiple IMIDs marked by pathogenic T cell activity (1, 2).

Several TNF/TNFR molecules are currently under investigation as therapeutic targets for IMIDs. Building on the success of large-molecule TNFi in the clinic, recent efforts have focused on the development of small-molecule TNFi designed for oral administration. Additionally, biologics targeting CD40L-CD40 and OX40L-OX40 interactions are in clinical development for IMIDs such as systemic lupus erythematosus, Sjögren’s syndrome, multiple sclerosis, atopic dermatitis, hidradenitis suppurativa, prurigo, and asthma (3). Anti-TL1A antibodies are in advanced clinical trials for inflammatory bowel diseases (IBD), with additional early-stage studies ongoing for atopic dermatitis, rheumatoid arthritis (RA), radiographic axial spondyloarthritis, hidradenitis suppurativa, systemic sclerosis-associated interstitial lung disease, and metabolic dysfunction-associated steatohepatitis (4). Antagonists targeting other TNF/TNFR superfamily members, including CD30L-CD30 (5, 6), GITRL/GITR (7), RANKL-RANK (8, 9), LIGHT-HVEM (10, 11), LTβ/LIGHT-LTβR (12, 13), and LTA (14, 15), have also been evaluated clinically as IMID therapy, though most failed to demonstrate sufficient efficacy to warrant additional studies.

Recent advances in immunotherapy have shifted from broad immunosuppression towards precise modulation of specific inflammatory pathways. Given the complexity of immune signaling, where redundant or synergistic crosstalk is common, targeting a single ligand-receptor interaction may be inadequate for refractory autoimmune and inflammatory conditions due to compensatory signaling. This limitation is evident from current treatments like TNFi that suffer from high primary non-response rates and loss of efficacy in initial responders, issues often managed through dose escalation, moving to a different TNFi, or switch to a different therapeutic class. Combination treatments that block redundant pathways could therefore provide a stronger and more durable suppression of inflammation, leading to improved disease control.

Synergistic interactions between TNF/TNFR signaling proteins have been documented across various preclinical models of inflammatory disease (16-23). Building upon these findings and the established role of these pathways in T-cell costimulation, we sought to systematically evaluate TNF/TNFR combinations as potential immunotolerance targets. We curated diverse datasets encompassing TNF/TNFR transcriptomics, genetics, druggability, genetic perturbation, inflammatory pathway regulation and literature mining scores, transformed the data using a desirability function, and derived a unified disease association score for each gene in RA and IBD, two IMIDs with substantial T cell involvement. Based on these disease association scores and the basal/activation-induced expression patterns in T cells, we selected ten TNF/TNFR targets (TNF, LTA, LTB, TL1A, OX40, CD40L, GITR, 4-1BB, CD30, and LIGHT) for combinatorial screening in mixed lymphocyte reactions using pharmacological inhibitors of their canonical interactions. Four combinations (co-inhibitions of TNF+CD40L, TNF+OX40L, CD40L+OX40L, and CD40L+LTβ/LIGHT) produced the most significant reduction in interleukin-2 (IL-2) and interferon gamma (IFN-γ) levels. RNA-Seq analysis revealed that several MLR activation-induced genes were downregulated exclusively in the combination groups, which suppressed T cell activation on a scale comparable to our positive control, CTLA4-Fc. Further, TNF+CD40L inhibition yielded the greatest suppression of pathways associated with T cell activation, proliferation, differentiation, and cytokine/chemokine production, indicating a broad dampening of alloreactive T cell response. Collectively, these findings support the potential of TNF/TNFR combination targeting as a tolerance-inducing therapeutic strategy with strong promise for further preclinical evaluation.

## Materials and methods

### Reagents

Adalimumab (Cat. HY-P9908), Pateclizumab (Cat. HY-P990034), Baminercept (Cat. HY-P99459), Amlitelimab (Cat. HY-P99434), Dapirolizumab (Cat. HY-P99842A), Quisovalimab (Cat. HY-P99810), Duvakitug (Cat. HY-P99842A), human IgG1 (Cat. HY-P99001) and human IgG4 (Cat. HY-P99003) were purchased from MedChemExpress. Human recombinant 4-1BB-Fc (Cat. 838-4B) was purchased from R&D Systems, and CTLA4-Fc (Cat. 310-05-500UG) was purchased from Peprotech. CD30Li (human α-CD30L) and GITRi (human α-GITR, hGITR-nc-Ab-#2) (24) are Pfizer Inc. internal tool compounds.

### Mixed lymphocyte reactions

Human PBMCs were enriched from leukopaks (StemCell Technologies, Cat. 70500.2) using Ficoll separation, following which CD3^+^ T cells and CD3^-^ cells were isolated using EasySep Human T Cell Isolation Kit (StemCell Technologies Cat. 17951) as per manufacturer’s recommendations. T cells were cultured in RPMI-1640 medium supplemented with 10% heat-inactivated fetal bovine serum, penicillin/streptomycin, and 0.05 mM 2-mercaptoethanol. CD3^-^cells were X-ray irradiated with 25 Gy (X-RAD320) and co-cultured 1:1 with CD3^+^ cells for 5 days in 96-well plates for a one-way allogeneic MLR in the presence of test agents. All molecules were used at 50 nM in the combination screen. 50 nM IgG1 and 50 nM CTLA4-Fc treatment groups served as control and comparator, respectively. Cell supernatant was collected on day 2 and day 5 for measurement of IL-2 and IFN-γ expression, respectively. Stimulator-responder pairs were prescreened for T cell activation in one-way MLR prior to use in experiments.

### Proliferation assay

For proliferation assays, the MLR was set-up with CD3^+^ cells that were stimulated with ImmunoCult T cell activator (Stemcell Technologies, Cat. 10991) for 3 days and rested for 2 days in fresh media. The rested cells were stained with CellTrace Violet (Invitrogen, Cat. C34557) and co-cultured with irradiated CD3^-^ cells for 5 days. On day 5, the cells were stained with CD3 antibody (Biolegend, Cat. 300440) and live/dead stain (Invitrogen, Cat. L34981) and assessed by flow cytometry to identify the percentage of CD3^+^ cells that have divided. Proliferation modeling done on FlowJo (v10, Waters Biosciences). Percent divided refers to the fraction of the original CD3^+^ population that has divided at least once, and expansion index is the overall increase (fold-increase) in CD3^+^ cell number.

### Cytokine profiling

Cytokines in the cell culture media were profiled using ELISA (Biolegend, Cat. 430115) for IFN-γ, and Meso Scale Discovery (MSD) for IL-2 (MesoScale, Cat. K151QQD-2) as per manufacturer’s assay protocol.

### Gene prioritization

Correlation analyses data were curated from non-cancerous tissue samples in the ArchS4 database queried using the correlationAnalyzeR package (v1.0.0) (25, 26). Cut-offs of Pearson correlation > 0.4, and adjusted *p*-value < 0.05 qualified genes for positive correlations with gene sets of interest and the percentage of positively correlated genes was the metric considered for the desirability transformation. Bulk transcriptomics data from RA, CD, and IBD patient diseased samples were curated from the gene expression omnibus (GEO) database and differential expression analyses done versus healthy tissue samples using Geo2R (27). Pseudo-bulk analyses of diseased RA (28), UC (29), and CD (30) scRNA-Seq samples were performed in R using custom scripts. For transcriptomics data, both the log fold changes in gene expression along with their adjusted *p*-values were considered for the desirability transformation. Text mining disease scores were obtained from the Open Targets platform (31). GWAS data on gene-disease associations were obtained from GWASCatalog (32). Target druggability and priority index scores were curated from Fang *et al.* (33). Pathway association scores were curated for the KEGG terms Inflammatory bowel disease (IBD), NF-kappa B signaling pathway, Rheumatoid arthritis, T cell receptor signaling pathway, and TNF signaling pathway from CytoReason. Genetic perturbation data on T cell activation was curated from Schmidt *et al.* (34). In vitro RNA-Seq data on human T cells stimulated with α-CD3/α-CD28 activating antibody and macrophages stimulated with LPS/IFN-γ were generated in-house. Each data type was transformed using desirability functions (35) that standardize heterogeneous evidence – such as *p*-values, FDR, effect sizes, fold changes, Priority index (33), and correlation metrics – into normalized scores suitable for weighted integration. This process yields an overall desirability score for each gene that represents the weight of evidence from Pfizer’s Gene Prioritization & Target Evaluation (GP&TE) integrated data for its involvement in the pathology of interest.

### RNA-Seq and analysis

To measure the expression of TNF/TNFR genes in T cells at baseline and with inflammatory activation, we performed RNA-Seq on CD3^+^ T cells from three donors of PBMCs before and after four hours of α-CD3/α-CD28 stimulation with ImmunoCult T cell activator (Stemcell Technologies, Cat. 10991). Cells were dissolved in RLT buffer (Qiagen, Cat. 79216) with 1% 2-mercaptoethanol for RNA isolation.

To assess the effects of select combinations of TNF/TNFR inhibitors on T cells activated in MLR, irradiated CD3^-^ cells were stained with CellTrace Violet (Invitrogen, Cat. C34557) prior to their co-culture with CD3^+^ in MLR in the presence of test agents. On day 5, the cells were harvested and stained with a live/dead stain (Invitrogen, Cat. L34981). The cells were then washed and resuspended in FACS buffer, and 500,000 live, CellTrace Violet negative cells were sorted per sample. The sorted cells were pelleted and dissolved in RLT buffer with 1% 2-mercaptoethanol.

RNA was isolated using NucleoSpin RNA Plus kits (Takara, Cat. 740984.50). Poly-A enrichment (stranded mRNA) library construction was performed, and an average of 30 million paired end reads were sequenced per sample. Adapters were trimmed using Cutadapt (v4.9), reads were aligned using STAR (v2.7.9a) using the GRCh38 assembly (release 111) and quantified using Salmon (v1.10.1). Downstream analysis was performed in R (v4.4.1). Genes were filtered using the filterByExpr function in EdgeR (v4.4.2), and voom function in Limma (v3.62.2) was used for normalization and differential gene expression analysis with a log-fold change of 1.5 and FDR-adjusted *P*-value of 0.05 being set as the threshold for determining differential expression. Gene Ontology (GO) enrichment analysis of the differentially expressed genes was conducted using clusterProfiler (v4.14.6) (36). Significance was defined by an FDR-adjusted *p*-value of less than 0.05.

### Statistical analysis

Statistical hypotheses testing was performed on Prism (v10, GraphPad software) or using custom R scripts.

## Results

### Computational gene prioritization of TNF/TNFR targets relevant to inflammatory disease and T cell activation

We used computational gene prioritization to assess the relevance of TNF/TNFR targets to inflammatory disease and T cell expression as a basis for a focused yet comprehensive combination screen *in vitro*. Disease association scoring for RA and IBD was performed in-house for the 47 TNF/TNFR genes using GP&TE framework, which is designed to prioritize genes for drug-target discovery by leveraging multi-omics data and providing an unbiased target evaluation based on integrated statistical analyses (**Fig. 1A**). GP&TE incorporates curated genetics, transcriptomics, text-mined literature associations, pathway regulation, druggability, and additional datasets into a unified model (**Supplemental Table S1, Fig. 1B**). The disease association scores we generated were comparable to RA and IBD association scores available on Open Targets and Data4Cure (**Supplemental Fig. S1**), and enabled a fine-grained ranking of TNF/TNFR genes that guided the choice of targets evaluated *in vitro* in combinations.

**Fig 1.**
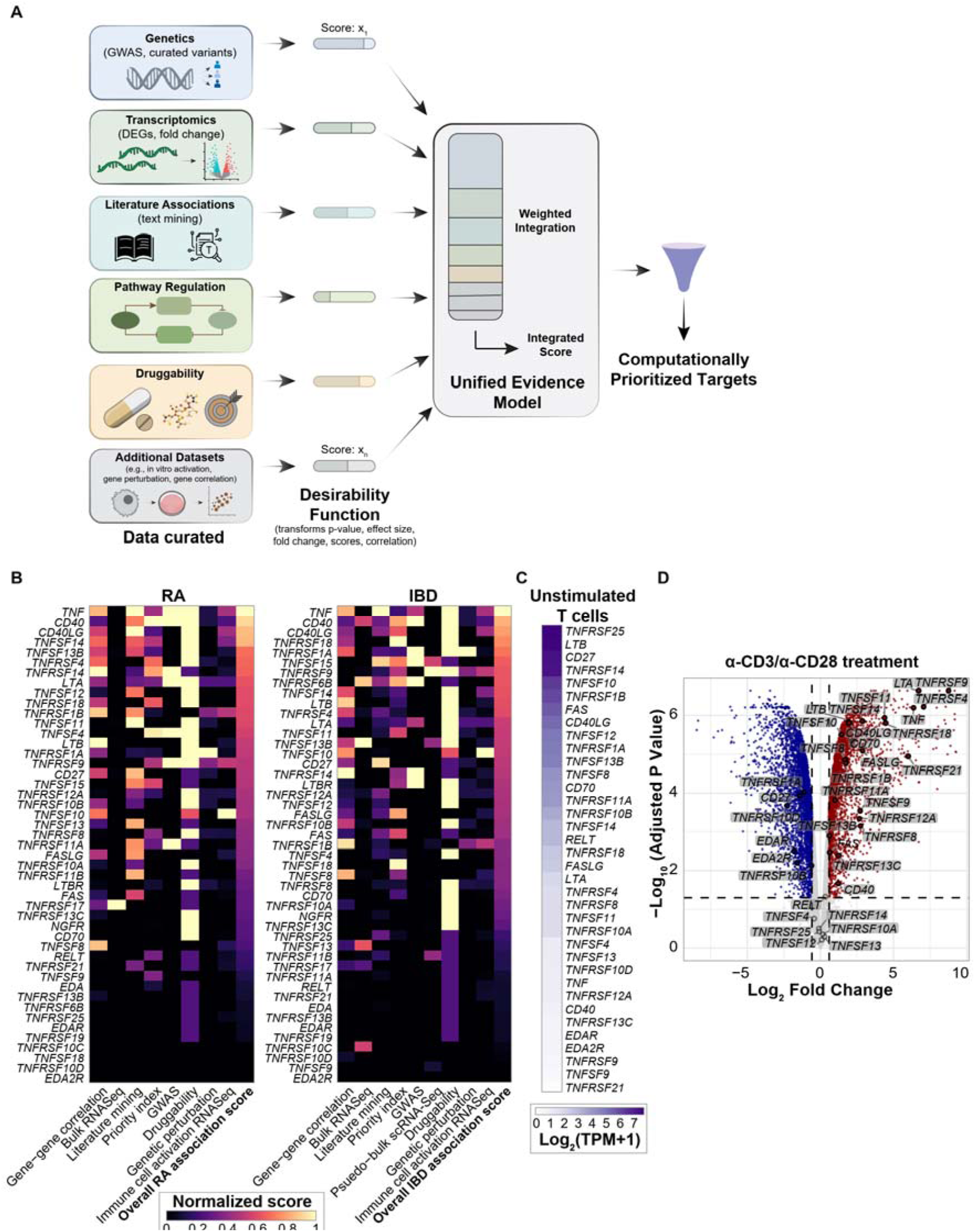
Computational gene prioritization reveals the TNF/TNFR targets relevant to inflammatory disease and T cell activation. (**A**) Schematic describing the data curation, transformation and integration steps undertaken to weigh the 47 TNF/TNFR genes. (**B**) RA and IBD gene scores for the TNF/TNFR genes ranked by overall disease association scores. Scores are normalized to the column maxima. (**C**) Expression of TNF/TNFR genes in unstimulated CD3^+^ human T cells. (**D**) Differential gene expression analysis of CD3^+^ T cells activated for 4 h using α-CD3/α-CD28 stimulus versus unstimulated cells. Data from three individual donors.

To further refine target selection, we complemented these computational findings with RNA-Seq data assessing TNF/TNFR family gene expression in naïve versus α-CD3/α-CD28-activated CD3^+^ T cells. TNFR molecules such as *TNFRSF14* (encoding HVEM), and *TNFRSF25* (encoding DR3), whose ligands are frequently upregulated in inflammatory diseases, were constitutively expressed in naïve T cells (**Fig. 1C**). Several prominent TNF/TNFR molecules, including *TNF*, *LTA*, *LTB*, *TNFRSF4* (encoding OX40), *CD40LG* (encoding CD40L), *TNFRSF8* (encoding CD30), *TNFRSF9* (encoding 4-1BB), and *TNFRSF18* (encoding GITR), were significantly upregulated upon T cell activation (**Fig. 1D**).

Based on computationally derived disease association scores and T cell expression profiles, we prioritized ten TNF–TNFR interactions inferred to be highly relevant to disease biology (Table 1, which summarizes the target selection rationale and expected effects of inhibition). These targets were subsequently evaluated by pharmacological blockade in the combination MLR screen.

**Table 1.**
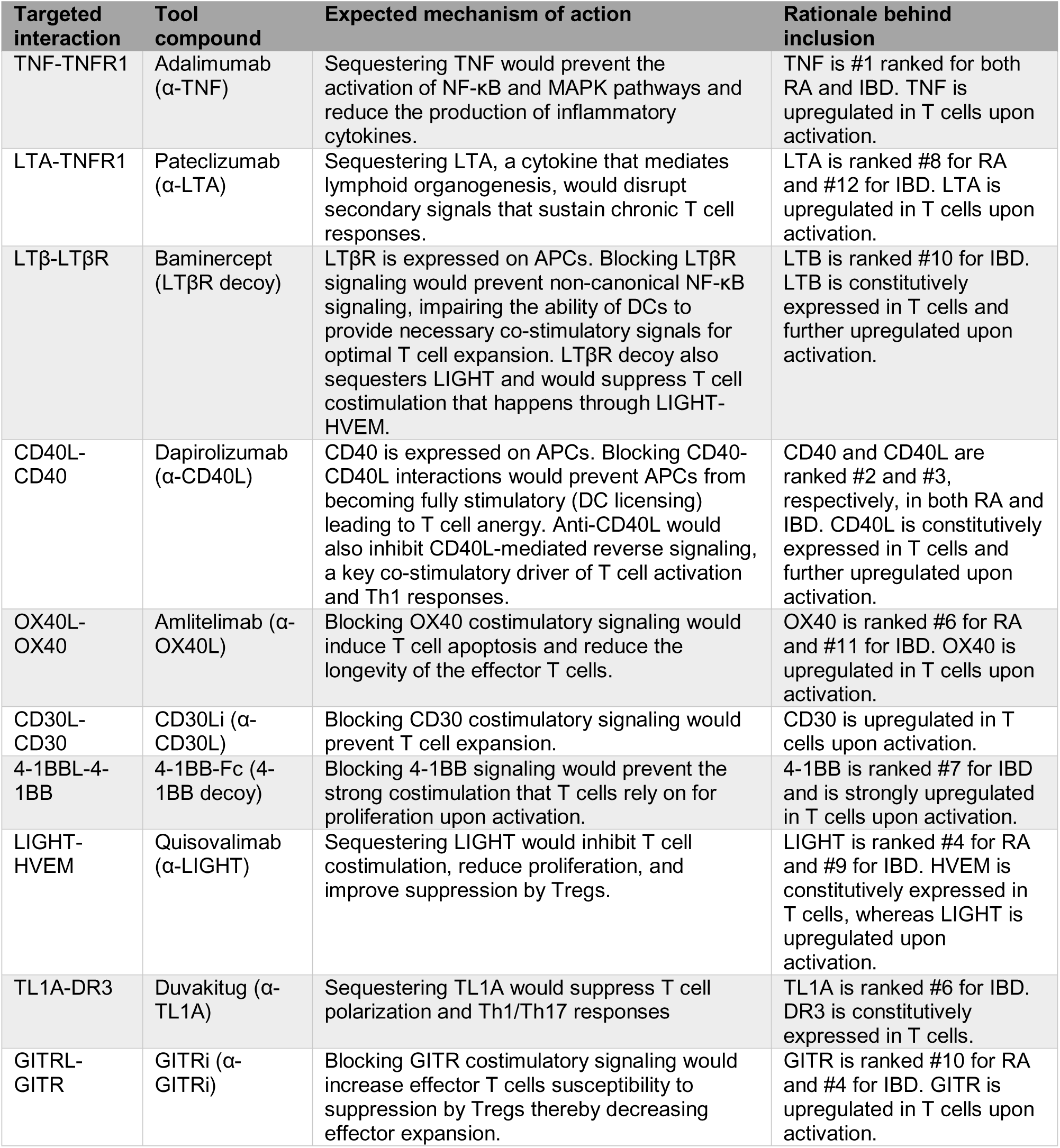
Computationally informed TNF/TNFR targets prioritized for in vitro combination screen in MLR.

| Targeted interaction | Tool compound | Expected mechanism of action | Rationale behind inclusion |
| --- | --- | --- | --- |
| TNF-TNFR1 | Adalimumab (α-TNF) | Sequestering TNF would prevent the activation of NF-κB and MAPK pathways and reduce the production of inflammatory cytokines. | TNF is #1 ranked for both RA and IBD. TNF is upregulated in T cells upon activation. |
| LTA-TNFR1 | Pateclizumab (α-LTA) | Sequestering LTA, a cytokine that mediates lymphoid organogenesis, would disrupt secondary signals that sustain chronic T cell responses. | LTA is ranked #8 for RA and #12 for IBD. LTA is upregulated in T cells upon activation. |
| LTβ-LTβR | Baminercept (LTβR decoy) | LTβR is expressed on APCs. Blocking LTβR signaling would prevent non-canonical NF-κB signaling, impairing the ability of DCs to provide necessary co-stimulatory signals for optimal T cell expansion. LTβR decoy also sequesters LIGHT and would suppress T cell costimulation that happens through LIGHT-HVEM. | LTB is ranked #10 for IBD. LTB is constitutively expressed in T cells and further upregulated upon activation. |
| CD40L-CD40 | Dapirolizumab (α-CD40L) | CD40 is expressed on APCs. Blocking CD40-CD40L interactions would prevent APCs from becoming fully stimulatory (DC licensing) leading to T cell anergy. Anti-CD40L would also inhibit CD40L-mediated reverse signaling, a key co-stimulatory driver of T cell activation and Th1 responses. | CD40 and CD40L are ranked #2 and #3, respectively, in both RA and IBD. CD40L is constitutively expressed in T cells and further upregulated upon activation. |
| OX40L-OX40 | Amlitelimab (α-OX40L) | Blocking OX40 costimulatory signaling would induce T cell apoptosis and reduce the longevity of the effector T cells. | OX40 is ranked #6 for RA and #11 for IBD. OX40 is upregulated in T cells upon activation. |
| CD30L-CD30 | CD30Li (α-CD30L) | Blocking CD30 costimulatory signaling would prevent T cell expansion. | CD30 is upregulated in T cells upon activation. |
| 4-1BBL-4-1BB | 4-1BB-Fc (4-1BB decoy) | Blocking 4-1BB signaling would prevent the strong costimulation that T cells rely on for proliferation upon activation. | 4-1BB is ranked #7 for IBD and is strongly upregulated in T cells upon activation. |
| LIGHT-HVEM | Quisovalimab (α-LIGHT) | Sequestering LIGHT would inhibit T cell costimulation, reduce proliferation, and improve suppression by Tregs. | LIGHT is ranked #4 for RA and #9 for IBD. HVEM is constitutively expressed in T cells, whereas LIGHT is upregulated upon activation. |
| TL1A-DR3 | Duvakitug (α-TL1A) | Sequestering TL1A would suppress T cell polarization and Th1/Th17 responses | TL1A is ranked #6 for IBD. DR3 is constitutively expressed in T cells. |
| GITRL-GITR | GITRi (α-GITRi) | Blocking GITR costimulatory signaling would increase effector T cells susceptibility to suppression by Tregs thereby decreasing effector expansion. | GITR is ranked #10 for RA and #4 for IBD. GITR is upregulated in T cells upon activation. |

### MLR Combination screen identifies four potent TNF/TNFR combinations that suppress T cell activation

We asked whether dual targeting of computationally-prioritized TNF/TNFR pathways could more effectively impose an immunotolerant state on T cells than single-pathway inhibition. For the shortlisted targets, we identified large-molecule pharmacological inhibitors that block their canonical interactions (**Table 1**). Seven inhibitors were commercially sourced clinical-stage molecules, two were internal preclinical tool compounds, and one (4-1BB-Fc) was a commercial decoy molecule. We performed allogeneic one-way MLR assays by coculturing CD3^+^ T cells with X-ray-irradiated CD3^-^ cells for 5 days and measured IL-2 and IFN-γ at days 2 and 5, respectively, in the presence of 50 nM of each test agents. Individual monotreatments were paired with 50 nM human IgG1. CTLA4-Fc was included as a positive comparator because it is known to effectively disrupt T cell costimulatory CD28 pathway and suppress MLR responses (37, 38). Among the monotreatments, Adalimumab (henceforth referred to as Ada, α-TNF), Dapirolizumab (Dapiro, α-CD40L), and Amlitelimab (Amlite, α-OX40L) significantly reduced IL-2 and IFN-γ levels relative to IgG1 control (**Fig. 2A**). Among the combinations, Ada+Dapiro yielded the highest suppression in IFN-γ and IL-2, reducing them to levels comparable to CTLA4-Fc (**Fig. 2B**). IFN-γ suppression by the Ada+Dapiro treatment was also significantly greater than with Ada or Dapiro alone. Dapiro in combination with Baminercept (Bam, LTβR decoy) also produced significant reductions in IL-2 and IFN-γ compared to IgG1 and exceeded both constituent monotreatments for IFN-γ inhibition. Ada+Amlite and Dapiro+Amlite were the other top-performing combinations, each causing significant reductions in IL-2 and IFN-γ compared with both IgG1 control and Amlite alone. Several additional combinations produced significant but more modest cytokine reductions relative to IgG1 (**Fig. S2**). Overall, Ada+Dapiro, Ada+Amlite, Dapiro+Amlite, and Dapiro+Bam were the four combinations that most robustly lowered proinflammatory cytokines in comparison to their respective monotreatments in this fixed-dose screen.

**Fig 2.**
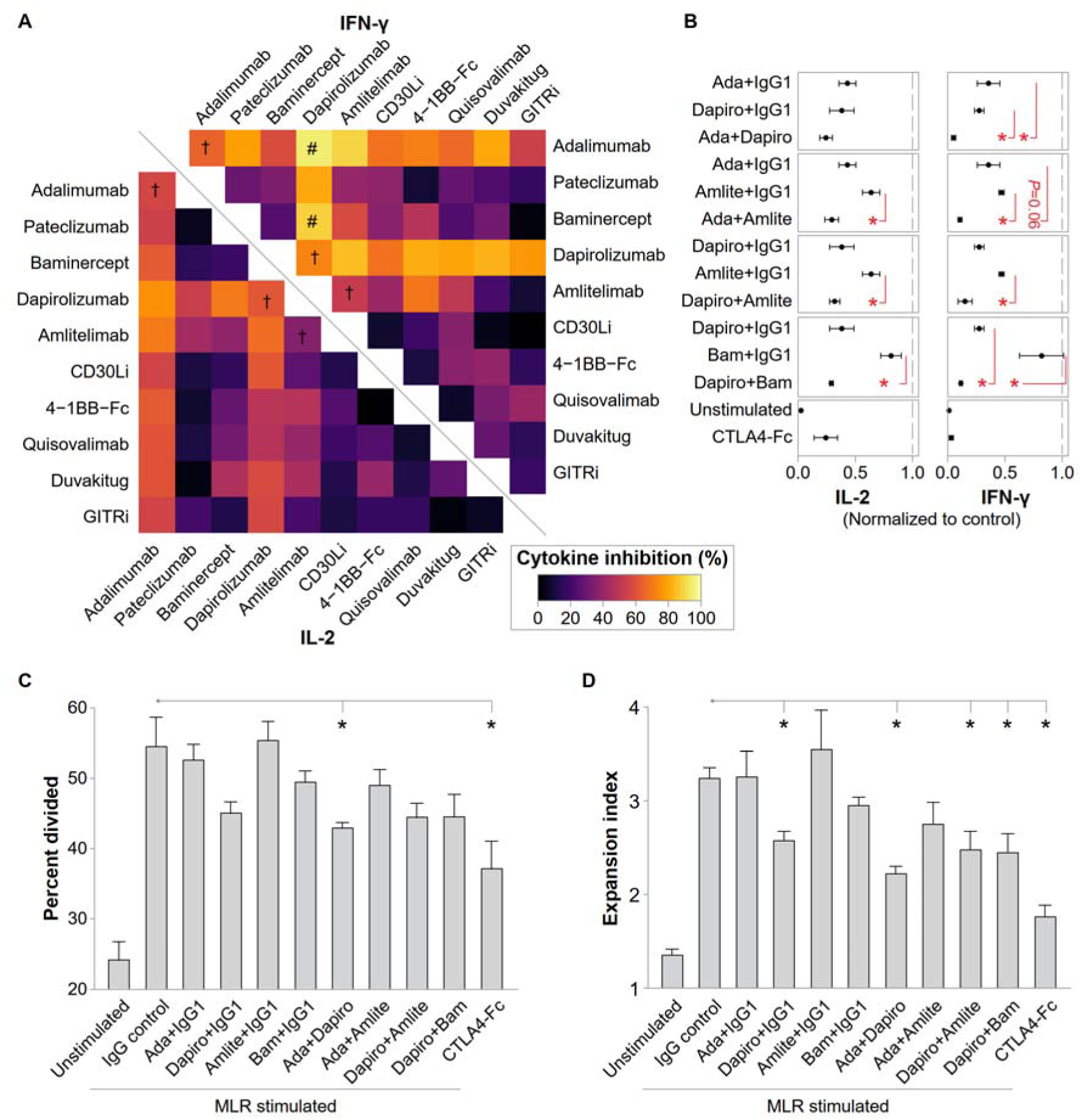
Combination screening of TNF/TNFR targets of interests in mixed lymphocyte reaction (MLR). (**A**) Interleukin-2 (IL-2) and interferon gamma (IFN-γ) inhibition in CD3^+^ cells during allogeneic one-way MLR measured at day 2 and day 5, respectively, treated with 50 nM combination pharmacological inhibitors of TNF/TNFR targets of interest. Monotreatment groups included 50 nM of human IgG1. Cytokine inhibition was calculated as percent reduction from IgG1 control treatment. Ill denotes *P*<0.05 in student t-test for monotreatments vs. IgG1 control, # denotes *P*<0.05 in student t-tests for combination treatment groups vs. IgG1 controls and both constituent monotreatments. (**B**) Normalized IL-2 and IFN-γ expression in four promising combinations of TNF/TNFR inhibitors. (**C-D**) Proliferation in CD3^+^ cells during allogeneic one-way MLR measured at day 5 in the presence of combination pharmacological inhibition of TNF/TNFR targets of interest.* denotes *P*<0.05 in student t-test. Data from three unique stimulator-responder MLR pairs, mean ± SEM plotted. Large-molecule pharmacological TNF/TNFR inhibitors used: Adalimumab (abbr. Ada, α-TNF), Pateclizumab (α-LTA), Baminercept (abbr. Bam, LTβR decoy), Dapirolizumab (abbr. Dapiro, α-CD40L), Amlitelimab (abbr. Amlite, α-OX40L), CD30Li (α-CD30L), 4-1BB-Fc (4-1BB decoy), Quisovalimab (α-LIGHT), Duvakitug (α-TL1A), and GITRi (α-GITR).

Given these results, we sought to examine T cell proliferation in the presence of these combinations. For the percentage of CD3^+^ cells that divided, Dapiro alone or in combination produced reductions that were not statistically significant, except for Ada+Dapiro which caused a significant decrease (**Fig. 2C**). Fold change in the numbers of CD3^+^ T cells, measured through expansion index, was significantly reduced in Dapiro monotreatment and combination treatment groups (**Fig. 2D**). Proliferation was reduced to similar levels by Dapiro alone and by the three combinations containing Dapiro for both proliferation readouts, indicating that CD40L-CD40 interactions between T cells and antigen-presenting cells (APCs) are critical drivers of T cell proliferation in the MLR setting.

We next conducted an equimolar combination dose-response MLR experiment as described above and measured IFN-γ at day 5 (**Fig. 3A-D**). Ada+Dapiro and Dapiro+Amlite reduced IFN-γ more effectively than their corresponding monotreatments, as reflected by lower IC_50_ and greater maximal suppression at their highest tested concentrations. Ada+Amlite and Dapiro+Bam exhibited slightly higher maximal effects than their corresponding monotreatments, although their IC_50_ levels were similar to Ada+IgG1 and Dapiro+IgG1, respectively. Collectively, these experiments indicate that the identified TNF/TNFR target combinations have the potential to enhance immunotolerance by dampening T cell activation and proliferation.

**Fig 3.**
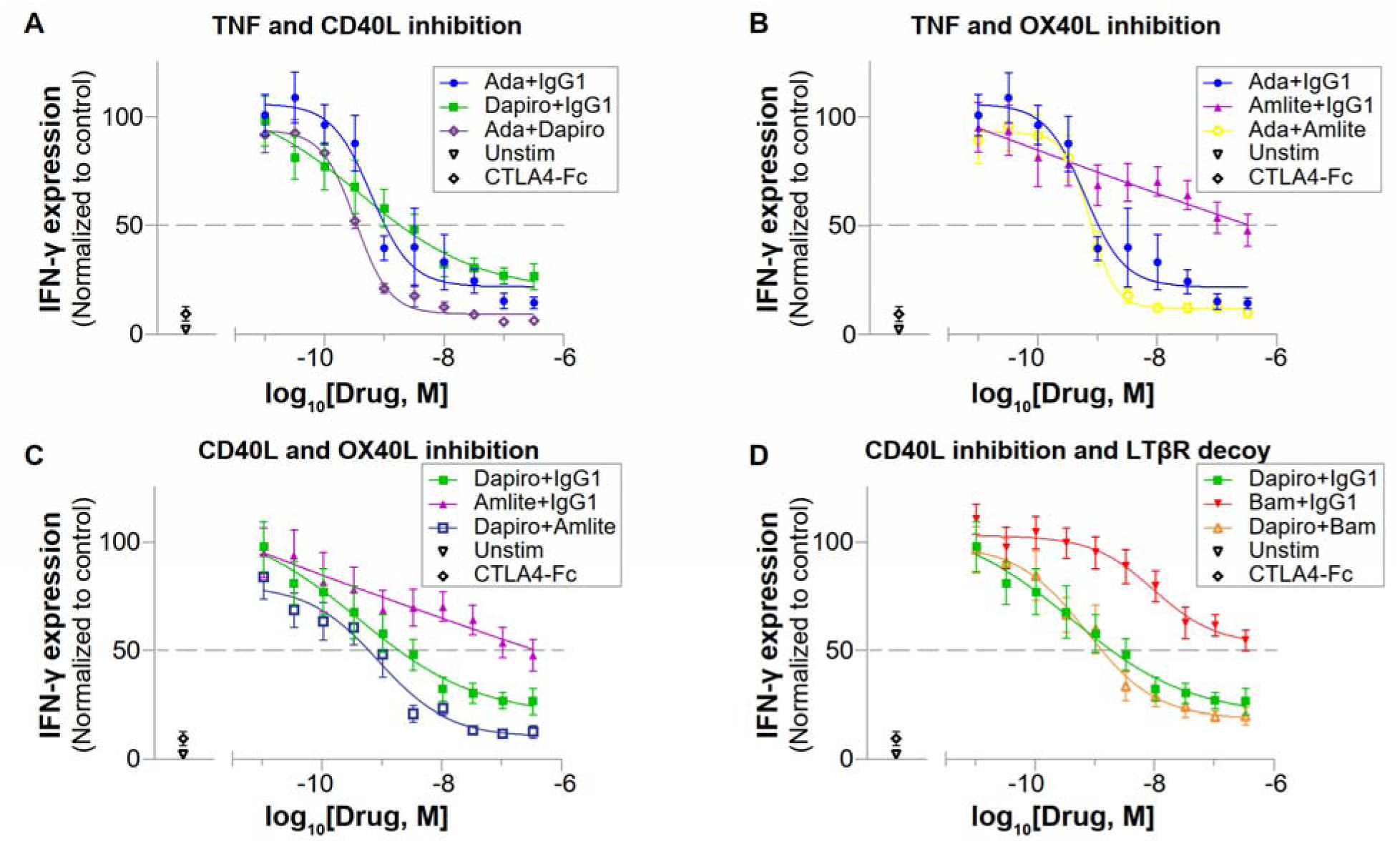
Four combinations of TNF/TNFR targets show promising T cell suppression. (**A-D**) IFN-γ expression in CD3^+^ cells during allogeneic one-way MLR measured at day 5 in the presence of equimolar combination of pharmacological inhibitors of TNF/TNFR targets of interest – (A) Ada and Dapiro, (B) Ada and Amlite, (C) Dapiro and Amlite, and (D) Dapiro and Bam. Data from four unique stimulator-responder pairs of MLR, mean ± SEM plotted. Large-molecule pharmacological TNF/TNFR inhibitors used: Adalimumab (abbr. Ada, α-TNF), Baminercept (abbr. Bam, LTβR decoy), Dapirolizumab (abbr. Dapiro, α-CD40L), and Amlitelimab (abbr. Amlite, α-OX40L).

### TNF/TNFR combinations downregulate MLR-upregulated genes in T cells

To dissect how these combinations remodel T cell activation states beyond cytokine production and proliferation, we characterized their transcriptional effects on T cells in MLR by RNA-Seq of flow sorted CD3^+^ cells harvested on day 5 of co-culture. 2×IgG1 and IgG1+IgG4 were used as negative controls (IgG4 included because Amlite is an IgG4 antibody) and CTLA4-Fc was used as a positive control. Additionally, unstimulated and α-CD3/α-CD28-stimulated T cells were included as inflammatory response controls. Compared with the no-drug MLR control, each monotreatment modulated fewer genes than the corresponding combination (**Fig. 4A**). Among monotreatments, Ada+IgG1 downregulated 122 genes, Ada+IgG4 160, Dapiro+IgG1 163, Dapiro+IgG4 131, Amlite+IgG1 66, and Bam+IgG1 3. Ada+Dapiro combination downregulated 524 genes, followed by Dapiro+Amlite (251), Dapiro+Bam (240), and Ada+Amlite (199), while CTLA4-Fc downregulated 387 genes. Analysis of the MLR-stimulated genes showed that the vast majority were modulated by combination treatments, whereas monotreatments induced more modest changes (**Fig. 4B**). Specifically, MLR-induced genes involved in cytotoxic effector function (*GZMA*, *GZMB*, *GZMH*, *GNLY*, *PRF1*), activation (*IL12RB2*, *STAP1*, *CD38*, *HOPX*), and trafficking (*CCR5*) were downregulated across all four combination treatments (**Fig. S3**).

**Fig 4.**
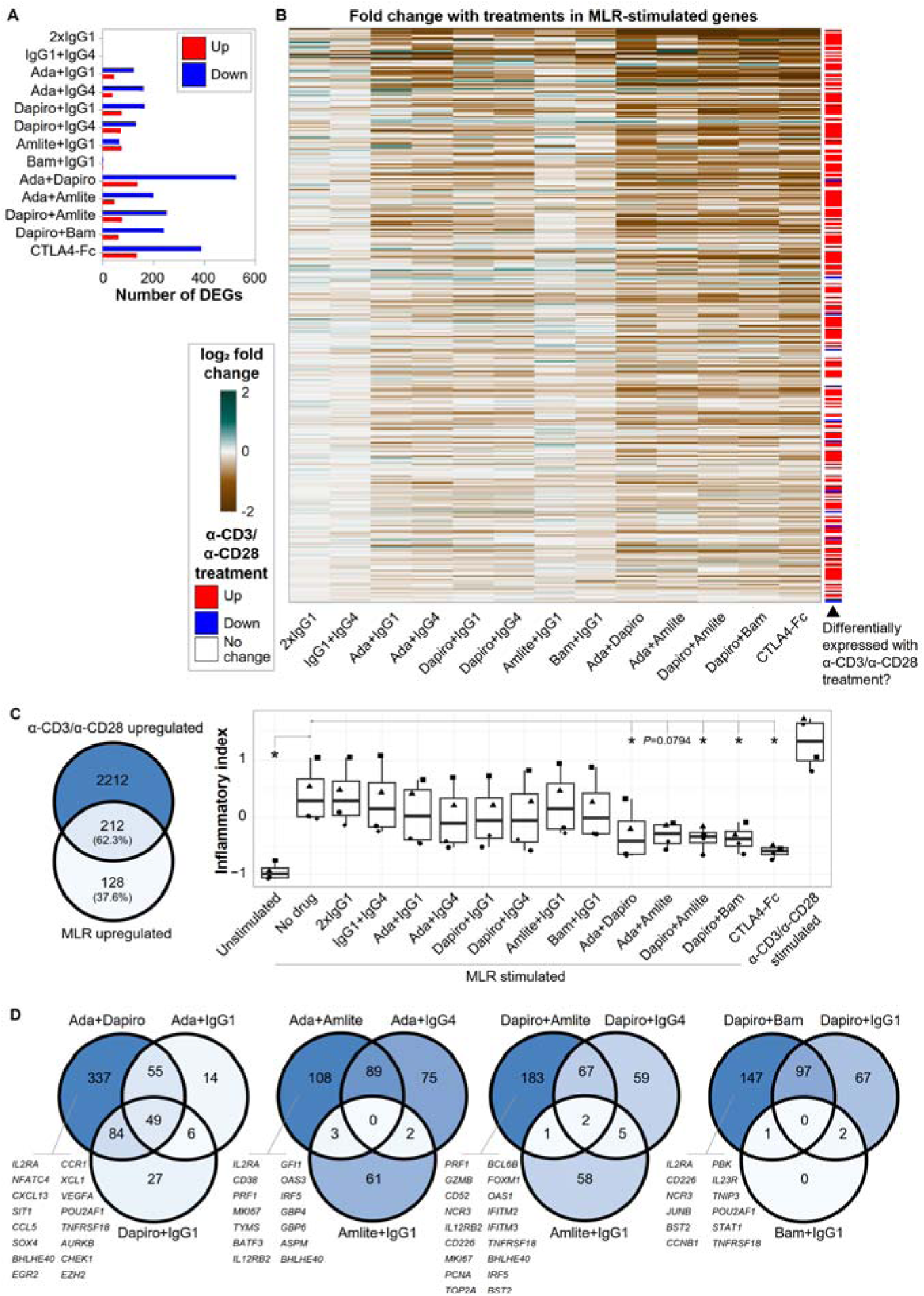
TNF/TNFR combination targets of interest downregulate MLR-upregulated genes in T cells. (**A**) Numbers of differentially expressed genes (DEGs) in response to the mono- and combination TNF/TNFR therapeutics of interest, versus the no drug treated MLR group (**B**) Fold change in MLR-upregulated genes with the addition of therapeutic agents. MLR- and α-CD3/α-CD28 upregulated genes were identified by comparing gene expression profiles of no drug MLR and α-CD3/α-CD28 treated cells, respectively, to that of unstimulated T cells. Genes are listed in the heatmap by fold change during MLR stimulation. (**C**) Inflammatory index composed of 212 genes commonly upregulated in both MLR- and α-CD3/α-CD28-stimulated cells. * denotes *P*<0.05 in Friedman test with Dunn’s multiple comparison post-hoc test. Individual points represent unique stimulator-responder MLR pairs. (**D**) Treatment-associated downregulated genes in each combination, compared to no drug treatment. RNA-Seq data from four unique stimulator-responder MLR pairs. Large-molecule pharmacological TNF/TNFR inhibitors used: Adalimumab (abbr. Ada, α-TNF), Baminercept (abbr. Bam, LTβR decoy), Dapirolizumab (abbr. Dapiro, α-CD40L), and Amlitelimab (abbr. Amlite, α-OX40L).

Additionally, genes associated with trafficking (*CXCR6*, *CCR2*), and immune checkpoints (*LAG3*, *HAVCR2* [encoding TIM-3]) that are upregulated upon T cell activation were selectively downregulated in the combinations involving Dapiro. Building an inflammatory index based on the 212 genes that are upregulated in T cells with both MLR- and α-CD3/α-CD28 stimulation (representing 62.3% of all MLR-upregulated genes), we observed significantly lower inflammatory index with the combination treatments Ada+Dapiro, Dapiro+Amlite, and Dapiro+Bam (**Fig. 4C**). Each of the four combinations also downregulated a substantial set of genes not affected by their constituent monotreatments (**Fig. 4D**). Ada+Dapiro uniquely downregulates 337 genes, representing 64.3% of the total genes downregulated by Ada+Dapiro, which were unaffected under Ada+IgG1 and Dapiro+IgG1 conditions. Similarly, Ada+Amlite, Dapiro+Amlite, and Dapiro+Bam uniquely downregulated 108 (54.2%), 183 (72.9%), and 147 (61.2%) genes, respectively. These combination-specific downregulated genes included key regulators of T cell activation (*IL2RA*, *NFATC4*, *TNFRSF18*, *STAT1*, *IL23R*) and proliferation (*MKI67*, *PCNA*, *TOP2A*). Gene ontology (GO) enrichment analysis of the genes downregulated by TNF/TNFR inhibitor combinations revealed significant enrichment for GO terms related to positive regulation of cytokine and chemokine production, immune activation, differentiation, and proliferation (**Fig. 5**). Ada alone and in combination (Ada+Dapiro and Ada+Amlite) downregulated gene sets that were particularly enriched for cytokine and chemokine related GO terms, whereas Dapiro alone and in combination (Ada+Dapiro, Dapiro+Amlite, and Dapiro+Bam) downregulated gene sets enriched for glycolysis. The gene set downregulated by Ada+Dapiro was particularly enriched for GO categories encompassing core T cell function including immune cell chemotaxis, production of type II interferons, IL-2, and IL-12, regulation of CD4^+^ αβ T cell activation, differentiation, and proliferation, and signaling via NF-κB, ERK1/2, and Ras. By comparison, gene sets downregulated by the other three combinations showed more modest enrichment in general T cell-related GO terms, indicating that Ada+Dapiro exerts the most pronounced suppressive effect on T cell activation among the four combinations. Collectively, these data suggest that dual TNF/TNFR blockade, particularly TNF+CD40L inhibition, enforces a broad attenuation of effector T cell programs characteristic of allogeneic activation.

**Fig 5.**
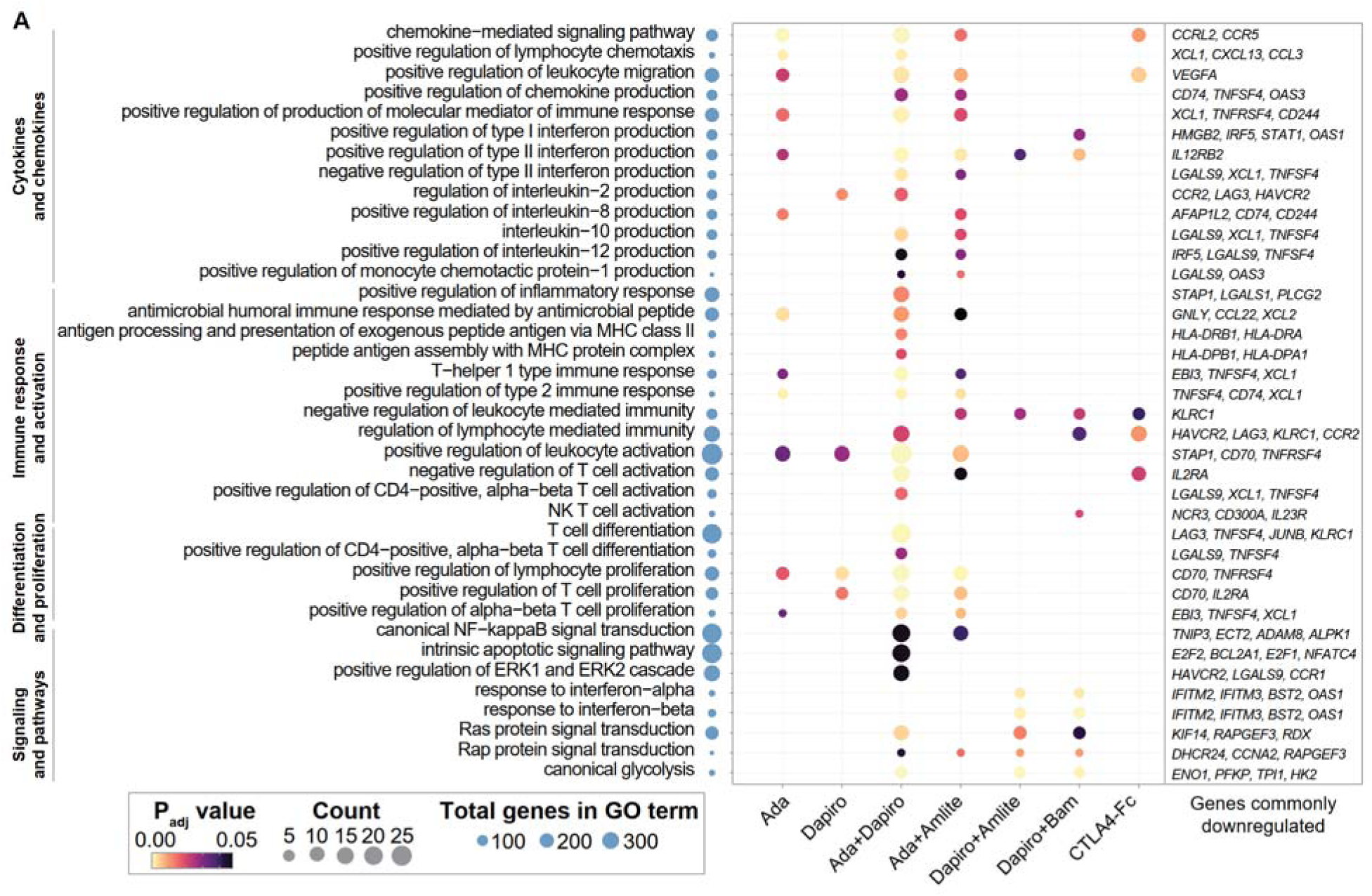
Gene ontology (GO) analysis of the genes downregulated with TNF/TNFR inhibitory treatment in comparison to no drug control in CD3^+^ cells during allogeneic one-way MLR showing representative GO terms spanning cytokines, chemotaxis, activation, differentiation, proliferation, and signaling pathways. GO terms associated with mitosis/general proliferation that were significant are not shown here. Large-molecule pharmacological TNF/TNFR inhibitors used: Adalimumab (abbr. Ada, α-TNF), Baminercept (abbr. Bam, LTβR decoy), Dapirolizumab (abbr. Dapiro, α-CD40L), and Amlitelimab (abbr. Amlite, α-OX40L).

## Discussion

Combination therapeutics within the TNF/TNFR superfamilies have garnered significant scientific interest due to their potential to overcome monotherapy resistance and address synergistic immune activation. Some of the synergistic signaling comes from the substantial cross-reactivity between ligands and receptors within these superfamilies. Furthermore, because several of these signaling proteins are upregulated following initial inflammatory activation, they can sustain inflammation by acting as redundant positive feedback cues. Consequently, several preclinical studies have explored how targeting these overlapping roles can provide superior therapeutic outcomes across diverse inflammatory conditions. CD30 and OX40 signaling drive allergic asthma in a murine house dust mite model where the dual blockade of OX40L and CD30L is required to suppress allergic airway inflammation and inhibit the effector memory T cell expansion (16). CD30 and OX40 signaling have also been observed to drive Th2 response in a mouse model of atopic dermatitis independently but combination antagonism failed to provide additional therapeutic benefit (17). TWEAK and TNF have been demonstrated to act synergistically in a imiquimod mouse model of psoriasis and promote disease progression by enhancing keratinocyte proliferation and inflammation and regulating an overlapping set of disease-signature genes (18). RANKL and TNF act synergistically in osteoclast activation; even low concentrations of TNF amplify osteoclastogenesis in response to sub-osteoclastogenic RANKL levels through positive feedback upregulation of RANKL (19, 20). Consistently, combined blockade of RANKL and TNF leads to greater protection against cartilage destruction than monotherapy in a TNF-driven polyarthritis mouse model (21). Similarly, TL1A and LIGHT signaling induce overlapping profibrotic programs in epithelial cells and fibroblasts, and dual inhibition results in significantly lower transforming growth factor β–driven fibroblast activation, collagen deposition, and smooth muscle accumulation in bleomycin-induced pulmonary fibrosis model compared with single treatments (22). Dual blockade of OX40 and TNF signaling have been demonstrated to significantly reduce proinflammatory cytokine production and CD4^+^ cell infiltration and improve histological score in a chronic colitis model of adoptive transfer of CD4^+^ CD45RB^high^ T cells (23). Taken together, these results suggest that the combined targeting of some TNF/TNFR proteins may be a viable strategy for IMID treatment.

Multiple TNF/TNFR combinations have been pursued as IMID therapeutics and advanced into clinical testing. Brivekimig is a TNF×OX40L bispecific nanobody currently in phase 2 clinical trials for hidradenitis suppurativa, ulcerative colitis, Crohn’s disease, and type 1 diabetes mellitus. In a humanized graft-versus-host disease (GvHD) mouse model, brivekimig significantly prolonged survival and reduced plasma IFN-γ levels compared with monospecific α-TNF or α-OX40L nanobodies (39). Brivekimig also showed promise in a phase 2a hidradenitis suppurativa study where the treatment group achieved a HiSCR50 clinical response rate of 67% versus 37% in the placebo group (NCT05849922, HS-OBTAIN) (40). Dual blockade of TNF and CD40/CD40L signaling has likewise attracted drug development efforts, including patents (41), and a clinical trial evaluating CD40L antagonism added to TNFi therapy in RA (NCT05306353). Given that TNFi treatment is less effective in IBD patients carrying a TL1A risk allele (42), combined TNF and TL1A blockade has also been explored. AMG-966, a human bispecific antibody targeting both TNF and TL1A, demonstrated encouraging preclinical activity for IBD. However, phase 1 clinical trials revealed a 98% rate of neutralizing anti-drug antibodies (ADAs) among participants leading to a loss of exposure (43), and the drug development was subsequently dropped.

Given the broad interest in combinations spanning distinct TNF/TNFR superfamily members, we systematically assessed the potential immunotolerance benefits achievable by combining targets most relevant in two model IMIDs, RA and IBD. We computationally prioritized ten TNF/TNFR interaction targets for combination screening in vitro. We used MLR as our screening assay given that it is among the most physiologically relevant in vitro systems to screen molecules belonging to the TNF/TNFR pathways. Notably, we did not observe any effect for the tested TNF/TNFR inhibitors in simpler experimental setups such as T cells stimulated with soluble α-CD3/α-CD28 antibody (data not shown), likely because T cells grown in isolation may not experience costimulatory TNF/TNFR signaling originating from APCs. Our MLR screen identified four combinations, TNF+CD40L, TNF+OX40L, CD40L+OX40L, and CD40L+LTβ, that produced the greatest inhibition of the inflammatory cytokines IL-2 and IFN-γ. Combinations involving CD40L-CD40 blockade also reduced T cell proliferation in MLR. RNA-Seq further confirmed the potential of these four combinations and established the TNF+CD40L inhibitory pair Ada+Dapiro as a particularly potent combination, with suppressive effects comparable to the positive control CTLA4-Fc. Notably, CTLA4-Fc acts at the very onset of T cell activation and is highly effective in MLR settings (44, 45), whereas the TNF/TNFR combinations we tested have the capacity to suppress inflammation in T cells already undergoing activation-induced changes. In acute GvHD, a condition closely related to the MLR model, the CTLA4 agonist Abatacept is used as prophylaxis. In contrast, TNFi such as infliximab and etanercept are used off-label as anti-inflammatory salvage therapy in steroid-refractory GvHD (46-48). By extension, GvHD patients whose disease escapes initial T cell costimulation blockade with Abatacept may also benefit from TNFi, highlighting the value of combination approaches for IMIDs that intervene at different stages of the immune response. Our results extend these prior efforts by using a data-driven framework to prioritize TNF/TNFR pairs and by demonstrating that selected combinations can rival CTLA4-Fc in suppressing human allogeneic T cell responses.

The targets we identified from our combination screens, TNF, CD40L-CD40, OX40L-OX40, and LTB/LIGHT, have the potential to act on distinct stages of the allogeneic T cell inflammatory response. Blockade of each of these targets has been shown either to suppress T cells in MLR or confer efficacy in GvHD models or trials. For example, TNF has broad effects on myeloid cells and lymphocytes, and functions as an early inflammatory cue that upregulates multiple costimulatory effector molecules. The TNFi treatments Ada and Infliximab reduce T cell proliferation in MLR by promoting regulatory macrophages (49). Similar to TNF, CD40L-CD40 signaling occurs early and supports immune synapse formation through dendritic cell (DC) licensing to activate antigen-specific T cells. This pathway drives the upregulation of major histocompatibility complex (MHC) molecules, and costimulatory receptors CD80/CD86, and augments APC production of TNF, IL-6, and IL-12, such that its blockade leads to T cells anergy (50). In addition, CD40L-CD40 inhibition may influence less well-characterized CD40L reverse signaling in T cells, mediated by receptor for activated C kinase 1 (RACK1) and implicated in IFN-γ-associated Th1 responses (51, 52). Letolizumab, an Fc-silent CD40L antibody, has been shown to be safe and to demonstrate preliminary efficacy in a phase I GvHD study when used prophylactically with standard of care (53). OX40-OX40L signaling is thought to occur later (at 24-48 h of T cell activation), where it prolongs T cell proliferation and survival, enhances IL-4 and IFN-γ production, and blocks suppression by regulatory T cells (Treg) (54-56). In MLR, Amlite reduces IL-2, TNF, and IL-4 production (57), while Telazorlimab (α-OX40) inhibits OX40L-driven T cell proliferation and NF-κB signaling (58). OX40 agonism on Tregs impairs their ability to suppress effector T cells in a skin allograft rejection mouse model (56), and intravenous immunoglobulin (IVIg) has been shown to deplete donor alloreactive CD4^+^ OX40^+^ cells coinciding with reduced proliferation and IFN-γ levels and increased survival in an acute GvHD rat model (59). In non-human primate GvHD models, Amlite combined with the mTOR inhibitor Sirolimus (a second-line therapeutic) achieved long-term GvHD-free survival and preserved Treg reconstitution (60). Soluble LTβR functions as a decoy receptor for LTαβ heterotrimers and LIGHT, inhibiting LTβ-mediated non-canonical NF-κB signaling and thereby limiting DC upregulation of CD80/CD86 and the provision of optimal co-stimulation for T cell expansion. LTβR decoys would also be expected to disrupt LIGHT-HVEM signaling, a costimulatory axis that supports robust activation and expansion of T cells, particularly CD8+ cytotoxic T lymphocytes. Soluble LTβR and HVEM have been shown to reduce T cell proliferation in MLR and to restrict anti-host cytotoxic T lymphocyte response in an acute GvHD mouse model (61). Interestingly, in the same model, soluble LTβR combined with α-CD40L enhanced survival and attenuated cytotoxic T lymphocyte responses by inducing donor T cell anergy to allogeneic antigens early during T cell activation (62). These previous studies underscore the significance of each of these promising TNF/TNFR targets in fostering immunotolerance. The presence of these intercellular signaling molecules in the synovium of RA (**Supplemental Fig. 4A**) and colonic lamina propria of IBD patients (**Supplemental Fig. 4B**) also highlights their functional involvement in disease tissue-specific pathology and potential as cross-indication therapeutic targets. Together, these results and those of our study underscore the therapeutic potential of combinatorial TNF/TNFR targeting to restore immune tolerance and support further preclinical development of such strategies for IMID treatment.

While our systematic approach successfully identified highly potent TNF/TNFR inhibitor combinations, this study has limitations inherent to the *in vitro* models employed. The allogeneic MLR is a robust and scalable system for evaluating T cell activation and costimulation, but it may not fully recapitulate the complex cytokine milieu, and cellular interactions present in disease conditions *in vivo*. Specifically, the MLR involving irradiated APCs may insufficiently model disease pathways driven by cytokines that are not extensively produced in this setting (63). For instance, the limited efficacy observed with TL1A combination blockade in our screen may reflect a lack of adequate TL1A production under these culture conditions, rather than the true absence of synergistic potential in clinical contexts like IBD where bacterial dysbiosis and the consequent myeloid cell response are connected to TL1A upregulation. Beyond these constraints, our initial screening approach utilized a fixed-dose strategy and focused primarily on circulating T cell responses, overlooking tissue-resident memory T cell dynamics and stromal cell interactions that are implicated in several IMIDs.

Future directions should focus on validating these top-performing combinations in more complex, disease-relevant preclinical models. Evaluating these combinations in humanized murine models of GvHD or tissue-specific inflammatory disease models would be essential to define their *in vivo* therapeutic window, assess safety, determine impact on Treg function and immune reconstitution, and allows for the investigation of the specific mechanisms by which dual blockade remodels the T cell landscape using tools like scRNA-Seq. Ultimately, the development of bispecific antibodies or optimized co-antibody formulations targeting these prioritized TNF/TNFR axes may offer a powerful therapeutic strategy to restore durable immunotolerance in patients with refractory IMIDs.

## Supporting information

Raw gene counts - T cell stimulation

Metadata - T cell stimulation

Raw gene counts - Mixed lymphocyte reactions

Metadata - Mixed lymphocyte reactions

## Disclosures

The authors are all current/past employees of Pfizer Inc. and may hold stock/options with Pfizer Inc.

## Data availability

The raw counts of RNA-Seq samples and their corresponding metadata have been made available as supplemental files.

## Acknowledgements

P.K.V. thanks Pfizer Inc.’s Postdoctoral Fellowship program for supporting and enabling their research activities. We would like to acknowledge the contributions of Vaughn Youngblood and Margery G.H. Pelletier for their expertise in flow cytometry and cell sorting for the RNA-Seq experiments, Ying Zhang for timely assistance with planning and execution of RNA-Seq experiments, and Heng Liu, Caitlyn Dickinson, Jason M. Jussif, and Wei Zheng for sharing their expertise on cellular irradiation, mixed lymphocyte reactions, and proliferation assays. We are also grateful for the research input from Brian Bates, Jason Arroyo, and Mike Primiano that helped shape this work.

## Author contributions

PKV: Conceptualization, Data curation, Formal analysis, Investigation, Methodology, Software, Validation, Visualization, Writing – original draft; WZ: Data curation, Formal analysis, Methodology, Software, Writing – review & editing; VM: Investigation, Methodology, Writing – review & editing; TAW: Conceptualization, Supervision, Writing – review & editing; JQ: Conceptualization, Project administration, Resources, Supervision, Writing – review & editing; FJK: Conceptualization, Project administration, Resources, Supervision, Writing – review & editing.

## Tables and Figures

**Table S1.** Multi-omics data curated on the TNF/TNFR superfamilies for gene prioritization, and subscores calculated for rheumatoid arthritis (RA) and inflammatory bowel diseases (IBD). [See file attached]

**Supplemental Fig S1.**
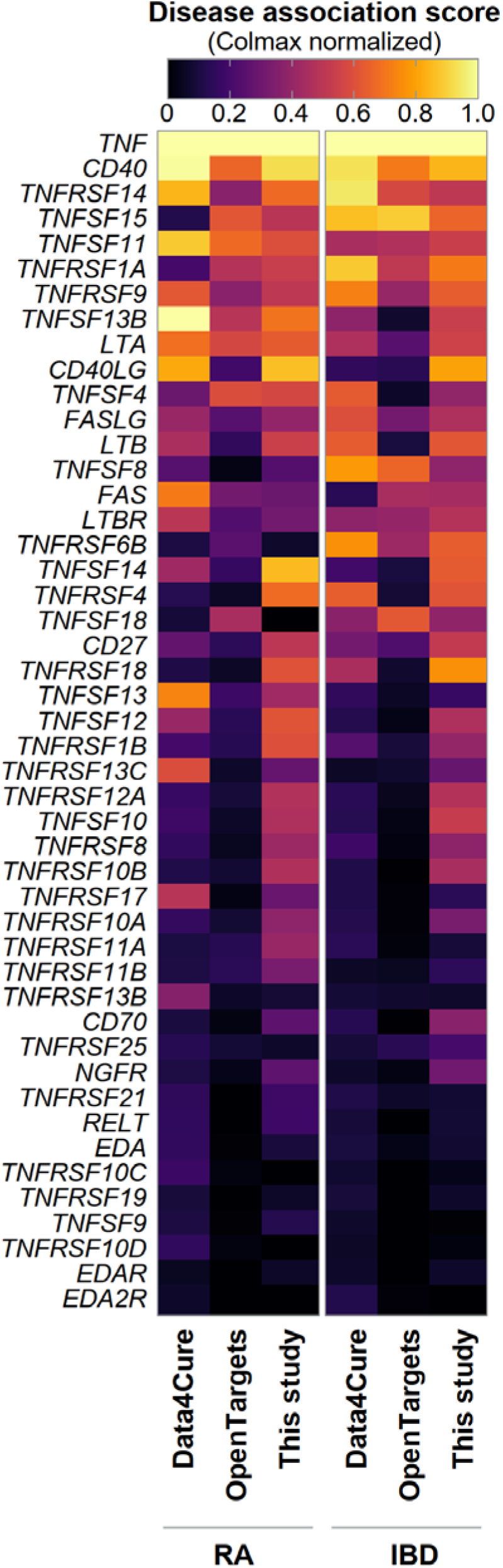
Comparison of disease scores from Data4Cure, Open Targets and GP&TE. Genes are ranked here by the average rank across all columns.

**Supplemental Fig S2.**
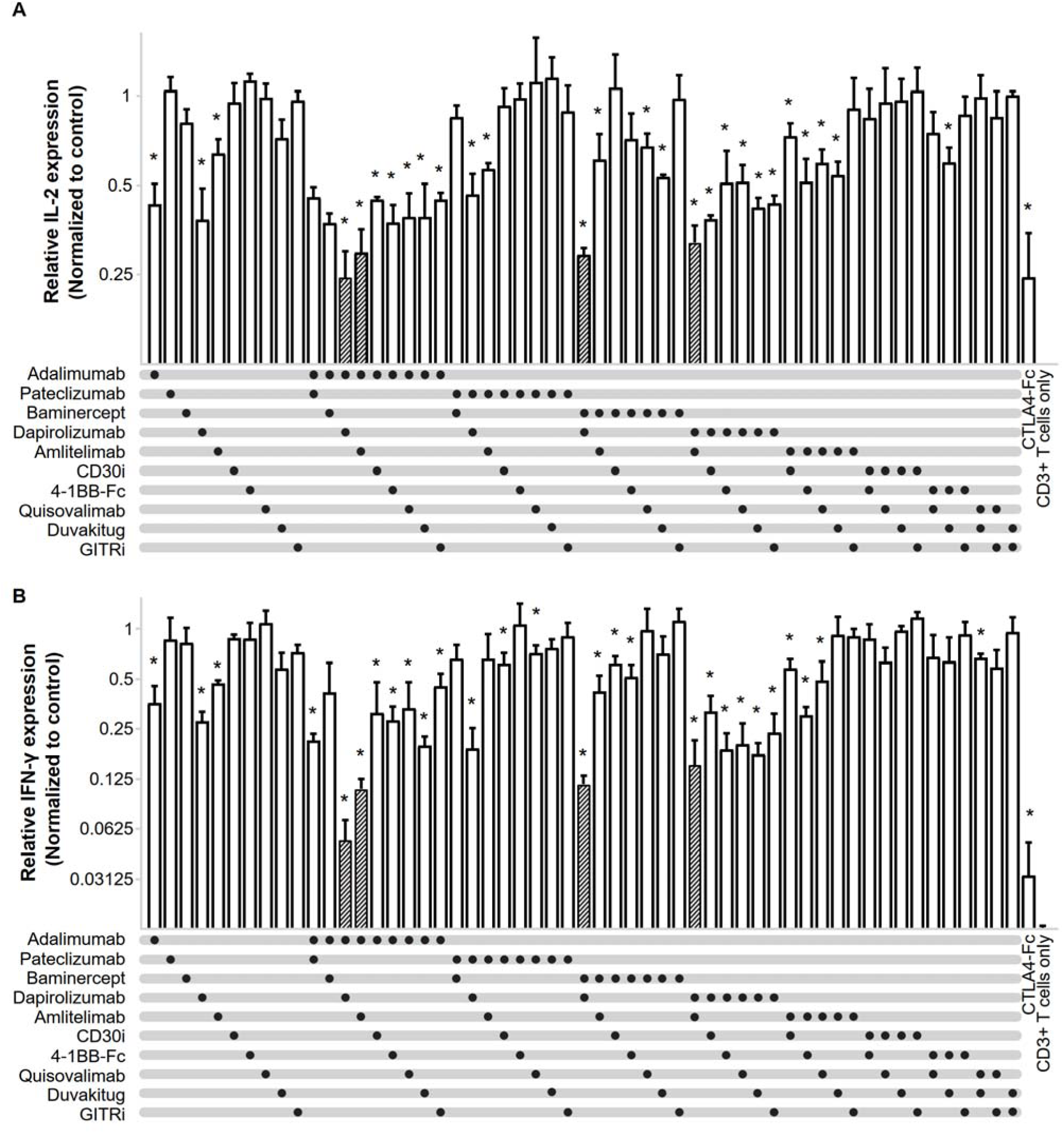
(**A-B**) Interleukin-2 (IL-2) and interferon gamma (IFN-γ) inhibition in CD3^+^ cells during allogeneic one-way MLR measured at day 2 and day 5, respectively, treated with 50 nM combination pharmacological inhibitors of TNF/TNFR targets of interest. Monotreatment groups included 50 nM of human IgG1. Cytokine expression normalized to IgG1 control treatment. * denotes *P*<0.05 in student t-test for monotreatments vs. IgG1 control. Data from three individual stimulator-responder MLR pairs, mean ± SEM plotted.

**Supplemental Fig S3.**
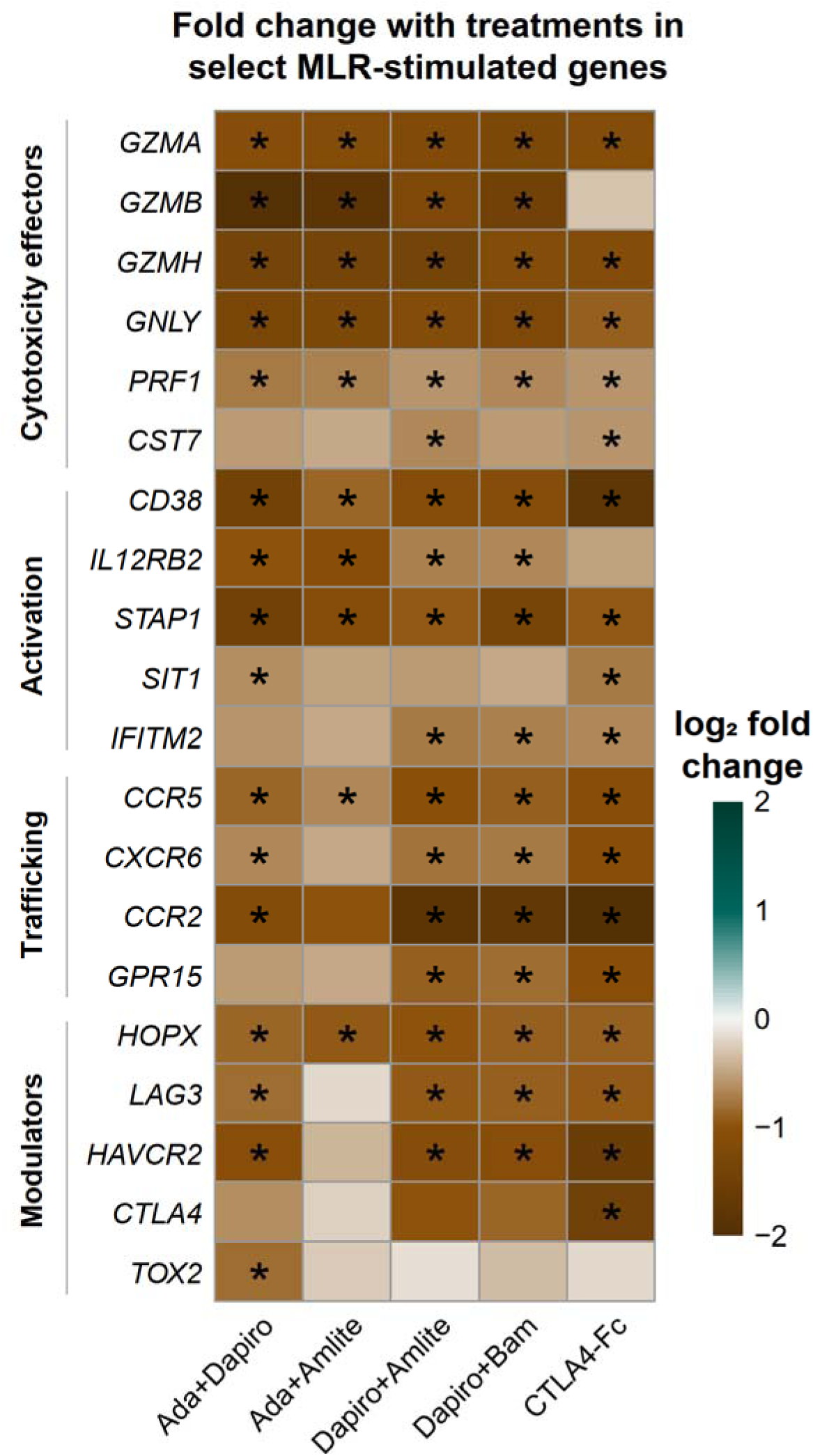
Fold change in MLR-upregulated genes with the addition of therapeutic combination agents, compared to CTLA4-Fc treatment. * denotes FDR-adjusted *P*<0.05 in DEG analysis.

**Supplemental Fig S4.**
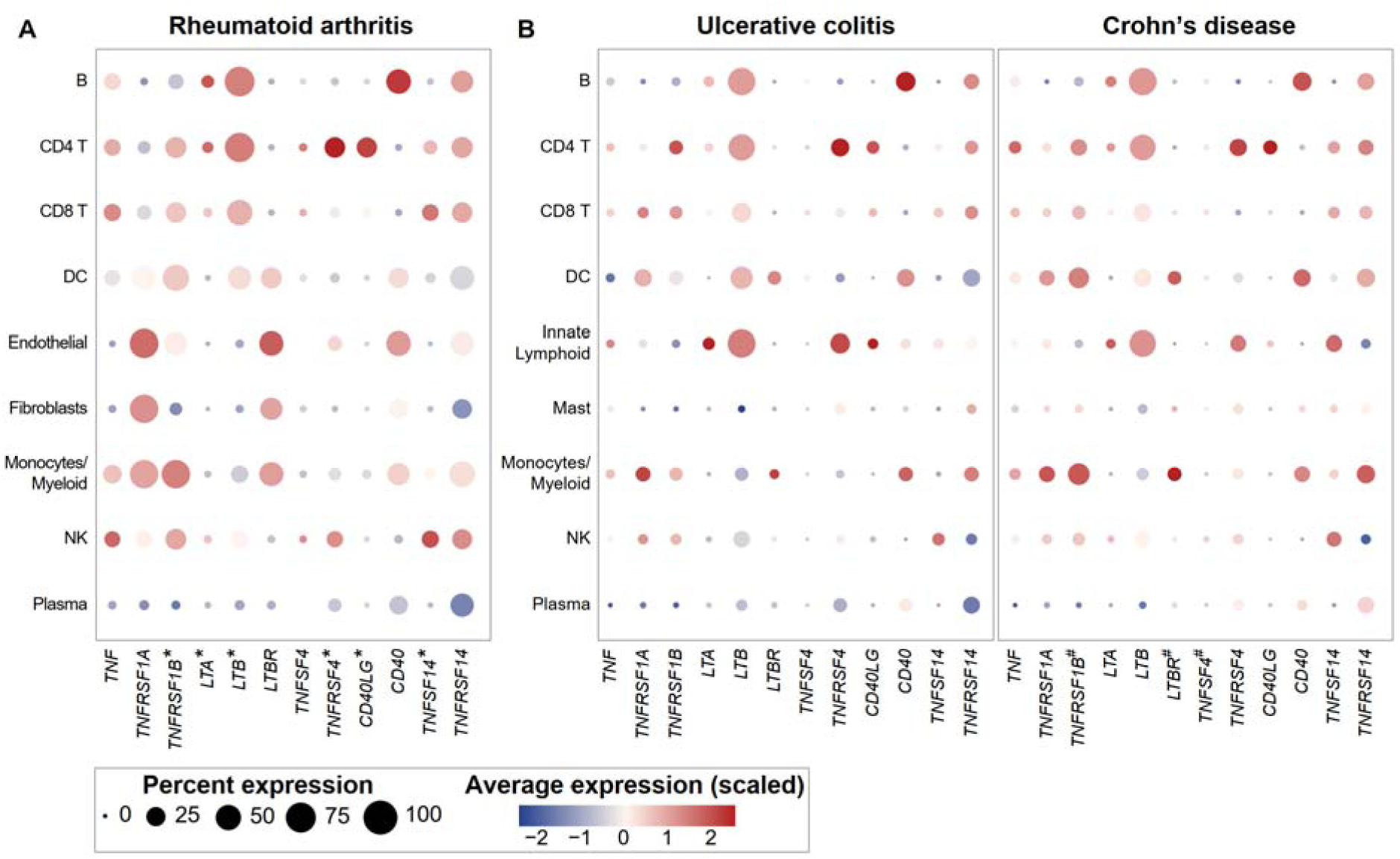
Expression of targets that are affected by the promising combination TNF/TNFR inhibitors in single cell RNA-Seq datasets derived from (**A**) synovial tissue of RA patients (28), and (**B**) inflamed colonic lamina propria of inflammatory IBD patients (29, 30). * and # denote significant upregulation (FDR-adjusted *P*<0.05 and fold-change > 1.5) in pseudo-bulk analyses comparing RA vs. osteoarthritis patients, and inflamed IBD vs. healthy subjects respectively.

## References

1. Watts TH, Yeung KKM, Yu T, Lee S, Eshraghisamani R. Tnf/Tnfr Superfamily Members in Costimulation of T Cell Responses-Revisited. Annu Rev Immunol (2025) 43(1):113–42. Epub 20250102. doi: 10.1146/annurev-immunol-082423-040557.

2. Croft M. The Role of Tnf Superfamily Members in T-Cell Function and Diseases. Nat Rev Immunol (2009) 9(4):271–85. doi: 10.1038/nri2526.

3. Veerasubramanian PK, Wynn TA, Quan J, Karlsson FJ. Targeting Tnf/Tnfr Superfamilies in Immune-Mediated Inflammatory Diseases. J Exp Med (2024) 221(11). Epub 20240919. doi: 10.1084/jem.20240806.

4. D’Amico F, Peyrin-Biroulet L, Jairath V, Danese S. Tl1a Blockade in Inflammatory Bowel Diseases: Clinical Trials to Watch. Med (2025) 6(12):100931. doi: 10.1016/j.medj.2025.100931.

5. Fecteau J-F, Renshaw M, Fransson J, Laurent O, Barnett B. Anti-Cd30l Antibodies and Uses Thereof. Google Patents (2024).

6. Fransson J, Connolly B, Dickerson CT, Fecteau J-F, Renshaw M, Laurent O, et al. Anti-Cd30l Antibodies, Formulations Therefor, and Uses Thereof. Google Patents (2025).

7. Verdino P, Cain PF, Lacerte MA, Lee SL, Wortinger MA. Gitr Antagonists and Methods of Using the Same. Google Patents (2024).

8. Cummings SR, San Martin J, McClung MR, Siris ES, Eastell R, Reid IR, et al. Denosumab for Prevention of Fractures in Postmenopausal Women with Osteoporosis. N Engl J Med (2009) 361(8):756–65. Epub 20090811. doi: 10.1056/NEJMoa0809493.

9. Cohen SB, Dore RK, Lane NE, Ory PA, Peterfy CG, Sharp JT, et al. Denosumab Treatment Effects on Structural Damage, Bone Mineral Density, and Bone Turnover in Rheumatoid Arthritis: A Twelve-Month, Multicenter, Randomized, Double-Blind, Placebo-Controlled, Phase Ii Clinical Trial. Arthritis Rheum (2008) 58(5):1299–309. doi: 10.1002/art.23417.

10. Perlin DS, Neil GA, Anderson C, Zafir-Lavie I, Raines S, Ware CF, et al. Randomized, Double-Blind, Controlled Trial of Human Anti-Light Monoclonal Antibody in Covid-19 Acute Respiratory Distress Syndrome. J Clin Invest (2022) 132(3). doi: 10.1172/jci153173.

11. Avalo Therapeutics Inc. Avalo Announces Topline Data from Phase 2 Peak Trial for Avtx-002 (Quisovalimab) in Patients with Non-Eosinophilic Asthma [(Press release)]. Avalo Therapeutics Inc. (2023) [updated 06-26-2023; cited 2023 12-31-2023]. Available from: https://ir.avalotx.com/press-releases/detail/173/avalo-announces-topline-data-from-phase-2-peak-trial-for.

12. Bienkowska J, Allaire N, Thai A, Goyal J, Plavina T, Nirula A, et al. Lymphotoxin-Light Pathway Regulates the Interferon Signature in Rheumatoid Arthritis. PLoS One (2014) 9(11):e112545. Epub 20141118. doi: 10.1371/journal.pone.0112545.

13. St.Clair EW, Baer AN, Wei C, Noaiseh G, Parke A, Coca A, et al. Clinical Efficacy and Safety of Baminercept, a Lymphotoxin Β Receptor Fusion Protein, in Primary Sjögren’s Syndrome. Arthritis & Rheumatology (2018) 70(9):1470–80. doi: 10.1002/art.40513.

14. Emu B, Luca D, Offutt C, Grogan JL, Rojkovich B, Williams MB, et al. Safety, Pharmacokinetics, and Biologic Activity of Pateclizumab, a Novel Monoclonal Antibody Targeting Lymphotoxin Α: Results of a Phase I Randomized, Placebo-Controlled Trial. Arthritis Res Ther (2012) 14(1):R6. Epub 20120108. doi: 10.1186/ar3554.

15. Kennedy WP, Simon JA, Offutt C, Horn P, Herman A, Townsend MJ, et al. Efficacy and Safety of Pateclizumab (Anti-Lymphotoxin-Α) Compared to Adalimumab in Rheumatoid Arthritis: A Head-to-Head Phase 2 Randomized Controlled Study (the Altara Study). Arthritis Res Ther (2014) 16(5):467. Epub 20141030. doi: 10.1186/s13075-014-0467-3.

16. Gracias DT, Sethi GS, Mehta AK, Miki H, Gupta RK, Yagita H, et al. Combination Blockade of Ox40l and Cd30l Inhibits Allergen-Driven Memory T(H)2 Cell Reactivity and Lung Inflammation. J Allergy Clin Immunol (2021) 147(6):2316–29. Epub 20201105. doi: 10.1016/j.jaci.2020.10.037.

17. Gupta RK, Figueroa DS, Ay F, Causton B, Abdollahi S, Croft M. Comparison of Cd30l and Ox40l Reveals Cd30l as a Promising Therapeutic Target in Atopic Dermatitis. Allergy (2025) 80(2):500–12. doi: 10.1111/all.16412.

18. Gupta RK, Gracias DT, Figueroa DS, Miki H, Miller J, Fung K, et al. Tweak Functions with Tnf and Il-17 on Keratinocytes and Is a Potential Target for Psoriasis Therapy. Science Immunology (2021) 6(65):eabi8823. doi: doi:10.1126/sciimmunol.abi8823.

19. Fuller K, Murphy C, Kirstein B, Fox SW, Chambers TJ. Tnfalpha Potently Activates Osteoclasts, through a Direct Action Independent of and Strongly Synergistic with Rankl. Endocrinology (2002) 143(3):1108–18. doi: 10.1210/endo.143.3.8701.

20. Lam J, Takeshita S, Barker JE, Kanagawa O, Ross FP, Teitelbaum SL. Tnf-Alpha Induces Osteoclastogenesis by Direct Stimulation of Macrophages Exposed to Permissive Levels of Rank Ligand. J Clin Invest (2000) 106(12):1481–8. doi: 10.1172/jci11176.

21. Zwerina J, Hayer S, Tohidast-Akrad M, Bergmeister H, Redlich K, Feige U, et al. Single and Combined Inhibition of Tumor Necrosis Factor, Interleukin-1, and Rankl Pathways in Tumor Necrosis Factor-Induced Arthritis: Effects on Synovial Inflammation, Bone Erosion, and Cartilage Destruction. Arthritis Rheum (2004) 50(1):277-90. doi: 10.1002/art.11487.

22. Steele H, Willicut A, Dell G, Ghastine A, Nguyen X-X, Lembicz P, et al. Combination Therapy Blocking Tnf Superfamily Members 14 and 15 Reverses Pulmonary Fibrosis. The Journal of Immunology (2025) 214(4):808–17. doi: 10.1093/jimmun/vkaf002.

23. Totsuka T, Kanai T, Uraushihara K, Iiyama R, Yamazaki M, Akiba H, et al. Therapeutic Effect of Anti-Ox40l and Anti-Tnf-Α Mabs in a Murine Model of Chronic Colitis. American Journal of Physiology-Gastrointestinal and Liver Physiology (2003) 284(4):G595–G603. doi: 10.1152/ajpgi.00450.2002.

24. Yan J, Min-DeBartolo J, Huang CS, Sharif MN, Li L, Fish S, et al. Ligand Non-Competitive Gitr Antibody Prevents Formation of the Obligatory Signal-Triggering Gitrl: Gitr Stoichiometry. Sci Rep (2025) 16(1):2752. Epub 20251218. doi: 10.1038/s41598-025-32541-6.

25. Lachmann A, Torre D, Keenan AB, Jagodnik KM, Lee HJ, Wang L, et al. Massive Mining of Publicly Available Rna-Seq Data from Human and Mouse. Nat Commun (2018) 9(1):1366. Epub 20180410. doi: 10.1038/s41467-018-03751-6.

26. Miller HE, Bishop AJR. Correlation Analyzer: Functional Predictions from Gene Co-Expression Correlations. BMC Bioinformatics (2021) 22(1):206. Epub 20210420. doi: 10.1186/s12859-021-04130-7.

27. Clough E, Barrett T, Wilhite SE, Ledoux P, Evangelista C, Kim IF, et al. Ncbi Geo: Archive for Gene Expression and Epigenomics Data Sets: 23-Year Update. Nucleic Acids Res (2024) 52(D1):D138-D44. doi: 10.1093/nar/gkad965.

28. Zhang F, Jonsson AH, Nathan A, Millard N, Curtis M, Xiao Q, et al. Deconstruction of Rheumatoid Arthritis Synovium Defines Inflammatory Subtypes. Nature (2023) 623(7987):616-24. Epub 20231108. doi: 10.1038/s41586-023-06708-y.

29. Smillie CS, Biton M, Ordovas-Montanes J, Sullivan KM, Burgin G, Graham DB, et al. Intra- and Inter-Cellular Rewiring of the Human Colon During Ulcerative Colitis. Cell (2019) 178(3):714–30 e22. doi: 10.1016/j.cell.2019.06.029.

30. Kong L, Pokatayev V, Lefkovith A, Carter GT, Creasey EA, Krishna C, et al. The Landscape of Immune Dysregulation in Crohn’s Disease Revealed through Single-Cell Transcriptomic Profiling in the Ileum and Colon. Immunity (2023) 56(12):2855. doi: 10.1016/j.immuni.2023.10.017.

31. Kafkas S, Dunham I, McEntyre J. Literature Evidence in Open Targets - a Target Validation Platform. J Biomed Semantics (2017) 8(1):20. Epub 20170606. doi: 10.1186/s13326-017-0131-3.

32. Sollis E, Mosaku A, Abid A, Buniello A, Cerezo M, Gil L, et al. The Nhgri-Ebi Gwas Catalog: Knowledgebase and Deposition Resource. Nucleic Acids Res (2023) 51(D1):D977–D85. doi: 10.1093/nar/gkac1010.

33. Fang H, De Wolf H, Knezevic B, Burnham KL, Osgood J, Sanniti A, et al. A Genetics-Led Approach Defines the Drug Target Landscape of 30 Immune-Related Traits. Nat Genet (2019) 51(7):1082–91. Epub 20190628. doi: 10.1038/s41588-019-0456-1.

34. Schmidt R, Steinhart Z, Layeghi M, Freimer JW, Bueno R, Nguyen VQ, et al. Crispr Activation and Interference Screens Decode Stimulation Responses in Primary Human T Cells. Science (2022) 375(6580):eabj4008. Epub 20220204. doi: 10.1126/science.abj4008.

35. Eidem HR, Steenwyk JL, Wisecaver JH, Capra JA, Abbot P, Rokas A. Integrate: A Desirability-Based Data Integration Framework for the Prioritization of Candidate Genes across Heterogeneous Omics and Its Application to Preterm Birth. BMC Med Genomics (2018) 11(1):107. Epub 20181119. doi: 10.1186/s12920-018-0426-y.

36. Wu T, Hu E, Xu S, Chen M, Guo P, Dai Z, et al. Clusterprofiler 4.0: A Universal Enrichment Tool for Interpreting Omics Data. The Innovation (2021) 2(3). doi: 10.1016/j.xinn.2021.100141.

37. Davis PM, Nadler SG, Rouleau KA, Suchard SJ. Abatacept (Ctla4-Ig) Modulates Human T-Cell Proliferation and Cytokine Production but Does Not Affect Lipopolysaccharide-Induced Tumor Necrosis Factor Alpha Production by Monocytes. Arthritis Research & Therapy (2005) 7(1):P21. doi: 10.1186/ar1542.

38. Levitsky J, Miller J, Huang X, Chandrasekaran D, Chen L, Mathew JM. Inhibitory Effects of Belatacept on Allospecific Regulatory T-Cell Generation in Humans. Transplantation (2013) 96(8):689–96. doi: 10.1097/TP.0b013e31829f1607.

39. Leeuw T, Šimaitė D, Heyninck K, Levin C, Cornelis S, Hijazi Y, et al. Combined Tnf-Α and Ox40l Targeting as a New Treatment Option for Hidradenitis Suppurativa. Journal of Allergy and Clinical Immunology: Global (2025) 4(3):100483. doi: 10.1016/j.jacig.2025.100483.

40. Sanofi. Press Release: Eadv: Sanofi’s Brivekimig Achieved Positive Results in Hidradenitis Suppurativa in Phase 2a Study. (2025).

41. Cha SH. Multispecific Antibodies, Compositions Comprising the Same, and Vectors and Uses Thereof. Google Patents (2021).

42. Endo K, Kakuta Y, Moroi R, Yamamoto K, Shiga H, Kuroha M, et al. Tl1a (Tnfsf15) Genotype Affects the Long-Term Therapeutic Outcomes of Anti-Tnfα Antibodies for Crohn’s Disease Patients. JGH Open (2020) 4(6):1108–13. Epub 20200801. doi: 10.1002/jgh3.12398.

43. Kroenke MA, Barger TE, Hu J, Miller MJ, Kalenian K, He L, et al. Immune Complex Formation Is Associated with Loss of Tolerance and an Antibody Response to Both Drug and Target. Front Immunol (2021) 12:782788. Epub 20211214. doi: 10.3389/fimmu.2021.782788.

44. Mayer E, Hölzl M, Ahmadi S, Dillinger B, Pilat N, Fuchs D, et al. Ctla4-Ig Immunosuppressive Activity at the Level of Dendritic Cell/T Cell Crosstalk. Int Immunopharmacol (2013) 15(3):638–45. Epub 20130220. doi: 10.1016/j.intimp.2013.02.007.

45. Davis PM, Nadler SG, Rouleau KA, Suchard SJ. Abatacept (Ctla4-Ig) Modulates Human T-Cell Proliferation and Cytokine Production but Does Not Affect Lipopolysaccharide-Induced Tumor Necrosis Factor Alpha Production by Monocytes. Arthritis Res Ther (2005) 7(Suppl 1):P21.

46. Patriarca F, Sperotto A, Damiani D, Morreale G, Bonifazi F, Olivieri A, et al. Infliximab Treatment for Steroid-Refractory Acute Graft-Versus-Host Disease. Haematologica (2004) 89(11):1352–9.

47. Couriel D, Saliba R, Hicks K, Ippoliti C, de Lima M, Hosing C, et al. Tumor Necrosis Factor-Alpha Blockade for the Treatment of Acute Gvhd. Blood (2004) 104(3):649–54. Epub 20040406. doi: 10.1182/blood-2003-12-4241.

48. Nygaard M, Andersen NS, Moser CE, Olesen G, Schjødt IM, Heilmann C, et al. Evaluation of Infliximab as Second-Line Treatment of Acute Graft Versus Host Disease - Validating Response on Day 7 and 28 as Predictors of Survival. Bone Marrow Transplant (2018) 53(7):844–51. Epub 20180201. doi: 10.1038/s41409-018-0099-3.

49. Vos ACW, Wildenberg ME, Duijvestein M, Verhaar AP, van den Brink GR, Hommes DW. Anti–Tumor Necrosis Factor-Α Antibodies Induce Regulatory Macrophages in an Fc Region-Dependent Manner. Gastroenterology (2011) 140(1):221–30.e3. doi: 10.1053/j.gastro.2010.10.008.

50. van Kooten C, Banchereau J. Cd40-Cd40 Ligand. Journal of leukocyte biology (2000) 67(1):2–17.

51. van Os BW, Vos WG, Bosmans LA, van Tiel CM, Toom Md, Beckers L, et al. Cd40l Modulates Cd4+ T-Cell Activation through Receptor for Activated C Kinase 1. European Journal of Immunology (2023) 53(12):2350520. doi: 10.1002/eji.202350520.

52. Blair PJ, Riley JL, Harlan DM, Abe R, Tadaki DK, Hoffmann SC, et al. Cd40 Ligand (Cd154) Triggers a Short-Term Cd4(+) T Cell Activation Response That Results in Secretion of Immunomodulatory Cytokines and Apoptosis. J Exp Med (2000) 191(4):651–60. doi: 10.1084/jem.191.4.651.

53. Khimani F, Ali H, Kim J, Cubitt C, Zhang S, Elmariah H, et al. Cd40 Ligand Blockade for Prevention of Graft-Versus-Host Disease. JCO Oncology Advances (2025) (2):e2400040. doi: 10.1200/oa-24-00040.

54. Gramaglia I, Weinberg AD, Lemon M, Croft M. Ox-40 Ligand: A Potent Costimulatory Molecule for Sustaining Primary Cd4 T Cell Responses. J Immunol (1998) 161(12):6510–7.

55. Croft M, So T, Duan W, Soroosh P. The Significance of Ox40 and Ox40l to T-Cell Biology and Immune Disease. Immunol Rev (2009) 229(1):173–91. doi: 10.1111/j.1600-065X.2009.00766.x.

56. Vu MD, Xiao X, Gao W, Degauque N, Chen M, Kroemer A, et al. Ox40 Costimulation Turns Off Foxp3+ Tregs. Blood (2007) 110(7):2501–10. doi: 10.1182/blood-2007-01-070748.

57. Saghari M, Gal P, Gilbert S, Yateman M, Porter-Brown B, Brennan N, et al. Ox40l Inhibition Suppresses Klh-Driven Immune Responses in Healthy Volunteers: A Randomized Controlled Trial Demonstrating Proof-of-Pharmacology for Ky1005. Clin Pharmacol Ther (2022) 111(5):1121–32. Epub 20220301. doi: 10.1002/cpt.2539.

58. Macoin J, Blein S, Monney T, Sancheti P, Reddy V, Back J, editors. Gbr830, a True Ox40 Antagonist Antibody with Potent Suppressive Effects on T Cell-Mediated Pathological Responses. ARTHRITIS & RHEUMATOLOGY; 2018: WILEY 111 RIVER ST, HOBOKEN 07030-5774, NJ USA.

59. Caccavelli L, Field AC, Betin V, Dreillard L, Belair MF, Bloch MF, et al. Normal Igg Protects against Acute Graft-Versus-Host Disease by Targeting Cd4(+)Cd134(+) Donor Alloreactive T Cells. Eur J Immunol (2001) 31(9):2781–90. doi: 10.1002/1521-4141(200109)31:9<2781::aid-immu2781>3.0.co;2-z.

60. Tkachev V, Furlan SN, Watkins B, Hunt DJ, Zheng HB, Panoskaltsis-Mortari A, et al. Combined Ox40l and Mtor Blockade Controls Effector T Cell Activation While Preserving T(Reg) Reconstitution after Transplant. Sci Transl Med (2017) 9(408). doi: 10.1126/scitranslmed.aan3085.

61. Tamada K, Shimozaki K, Chapoval AI, Zhai Y, Su J, Chen S-F, et al. Light, a Tnf-Like Molecule, Costimulates T Cell Proliferation and Is Required for Dendritic Cell-Mediated Allogeneic T Cell Response1. The Journal of Immunology (2000) 164(8):4105–10. doi: 10.4049/jimmunol.164.8.4105.

62. Tamada K, Tamura H, Flies D, Fu Y-X, Celis E, Pease LR, et al. Blockade of Light/Ltβ and Cd40 Signaling Induces Allospecific T Cell Anergy, Preventing Graft-Versus-Host Disease. The Journal of Clinical Investigation (2002) 109(4):549–57. doi: 10.1172/JCI13604.

63. Merrick A, Errington F, Milward K, O’Donnell D, Harrington K, Bateman A, et al. Immunosuppressive Effects of Radiation on Human Dendritic Cells: Reduced Il-12 Production on Activation and Impairment of Naïve T-Cell Priming. British Journal of Cancer (2005) 92(8):1450–8. doi: 10.1038/sj.bjc.6602518.

